# Divisively normalized integration of multisensory error information develops motor memories specific to vision and proprioception

**DOI:** 10.1101/561332

**Authors:** Takuji Hayashi, Yutaro Kato, Daichi Nozaki

**Affiliations:** Graduate School of Engineering, Tokyo University of Agriculture and Technology, Tokyo, Japan; School of Engineering and Applied Sciences, Harvard University, Cambridge, Massachusetts, USA; Japan Society for the Promotion of Science, Tokyo, Japan; Graduate School of Education, The University of Tokyo, Tokyo, Japan

## Abstract

Both visual and proprioceptive information contribute to accurate limb movement, but the mechanism of integration of these different modality signals for movement control and learning remains controversial. Here, we aimed to elucidate the mechanism of integration by examining motor adaptation when various combinations of visual and proprioceptive perturbations were applied during reaching movements. We found that the movement corrections by adaptation were explained by a mechanism known as a divisive normalization, which was previously reported to be the mechanism underlying the integration of multisensory signals in neurons. Furthermore, we found evidence that the motor memory for each sensory modality was formed separately and the outputs from these memories were integrated. These results provide a novel view of the utilization of different sensory modality signals in motor control and adaptation.

## INTRODUCTION

When we perform a motor skill, we rely on multiple sensory information, primarily vision and proprioception. Using a golf approach shot as an example, the visual system offers information regarding the pin, ball, and obstacles (e.g., bunkers or water hazards), while proprioceptive feedback offers information regarding the limbs and club. If we fail to accomplish the desired shot, the sensorimotor system corrects the subsequent shot based on the error between the actual and predicted sensory feedback, i.e., sensory prediction error^1^. Experimental paradigms with visuomotor rotation^2^ or force field perturbation^3,4^ have demonstrated the contribution of both modalities of sensory prediction errors in motor adaptation.

Importantly, the motor adaptation system receives sensory information from both modalities simultaneously. The contribution of the information provided by each sensory modality has been investigated by eliminating visual information^5^, recruiting blind^6^ and deafferent subjects^7,8^, or disturbing proprioceptive information^9^. However, the computational principle of the utilization of the sensory information in the motor adaptation system has not been fully elucidated. Previous studies have reported that sensory signals in different modalities are Bayesian-optimally integrated to estimate the size of object (haptic and vision^10^) and to locate the perceived position of a limb (proprioception and vision^11,12^). A straightforward solution would be that the error is calculated by a similar optimal integration mechanism for motor adaptation. However, such correspondence between the perception of error size and the error information processing for motor adaptation is not necessarily guaranteed, because motor adaptation could progress without awareness of movement error (i.e., implicitly), as was typically observed when the perturbation size was gradually increased^13–16^.

Indeed, this mechanism cannot explain the experimental result when various amounts of errors are imposed during a reaching movement. Linear summation models including the optimal integration model^10–12^ predict that the aftereffect, which reflects the size of the integrated error, should linearly increase with the error size. However, the aftereffect does not increase with the size of an error, but instead saturates^17–19^. In order to explain such a saturation effect, Wei and Kording^17^ proposed that the greater dissociation between visual and proprioceptive information reduces the relevance of the error, which results in decreasing the gain of adaptation. In contrast, Marko et al.^19^ from their investigation on how the aftereffects are modulated with the sizes of visual and proprioceptive errors, concluded that visual and proprioceptive errors independently contributed to the aftereffect. However, an additional mechanism was needed to explain the reduction in the adaptation gain with error size, which they considered as inherent characteristics of the motor adaptation system. Critically, they also demonstrated that even when the congruence between visual and proprioceptive information was maintained, the saturation of aftereffect with the error size was still present, which was inconsistent with the notion suggested by Wei and Kording^17^. As exemplified by this inconsistency, we have not reached a coherent understanding of how sensory prediction error is calculated from errors provided in different modalities.

More recently, another possible idea, divisive normalization, has been proposed as a mechanism of multisensory integration. This idea suggests that the neuronal activity is normalized by the pooled activities of neurons and is considered to be a canonical neural computation in the brain^20^. Ohshiro et al.^21,22^ have demonstrated that the multimodal (visual and vestibular) neuronal activity pattern can be explained by the divisive normalization mechanism: The stimulus for one of the modalities, even if it did not elicit any response in the multisensory neurons, could suppress the response to the stimulus for another modality when both stimuli were simultaneously presented (cross-modal suppression).

If the neuronal circuit for motor adaptation has similar characteristics, it is possible that the motor adaptation pattern induced by various combinations of visual and proprioceptive errors follows the pattern predicted by the divisive normalization mechanism. In this study, we explore this possibility that no previous studies have never examined. Specifically, we considered the following divisive normalization model for motor adaptation^23,24^:

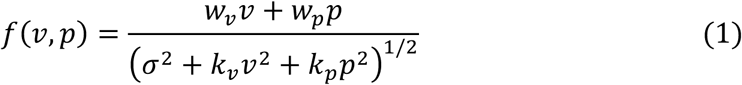

where *f*(*v*, *p*) is the aftereffect, *v* and *p* are the visual and proprioceptive error (or perturbation) imposed during a reaching movement, respectively, and *w*_*v*_, *w*_*p*_, *k*_*v*_, *k*_*p*_, and *σ* are constants (It should be noted that *σ* can be set to unity without loss of generality).

In order to determine the shape of *f*(*v*, *p*) experimentally, we need to precisely impose various combinations of visual and proprioceptive errors with wide range of magnitudes. The conventional experiments using force field and visual rotation are not appropriate for this purpose because the size of error should randomly vary among trials. Thus, we attempted to use the force channel method^25^ to impose visual and proprioceptive errors precisely and independently by deviating the visual cursor and hand movement directions from a target direction. In the subsequent probe trial, the force channel method was also used to measure the aftereffects to quantify the single-trial motor adaptation. The experimental results demonstrate that the aftereffects reasonably followed the pattern predicted by the divisive normalization mechanism (Eq. 1).

Another notable issue is at which stage the integration of visual and proprioceptive information occurs. The straightforward interpretation is that a motor memory is updated by the integrated error information^19^. However, there is an alternative interpretation that each modality has its own motor memory^26^ updated by divisively normalized error and the outputs from both motor memories are integrated. Our experiment supports the second interpretation that visual and proprioceptive memories are separately created and integrated at the output stage. The divisively normalized integration mechanism proposed in the present study provides novel insight into the integration of visual and proprioceptive information for motor adaptation.

## RESULTS

We sought to investigate how the aftereffects were induced by combinations of perturbations for vision and proprioception. When participants performed 10 cm reaching movements towards a front target, the cursor and hand movement trajectories were independently perturbed toward different directions with a force channel^25^ (Fig. 1a). In total, there were 35 combinations of visual perturbations (Fig. 1a red, 7 patterns of cursor directions, 0°, ± 15°, ± 30°, and ± 45°) and proprioceptive perturbations (Fig. 1a blue, 5 patterns of hand directions, 0°, ± 15°, and ± 30°). In the subsequent probe trial toward the front target, the force channel trial was also used to evaluate the aftereffect by measuring the lateral force against the channel. After the probe trial, 2 null trials (without the force channel) were performed to washout the possible adaptation effects (Fig. 1b). The cursor was continuously visible during movements in Experiment 1 (Online feedback condition, Fig. 1c), while the cursor appeared only after the movement completion in Experiment 2 (Endpoint feedback condition, Fig. 1d).

**Fig. 1.**
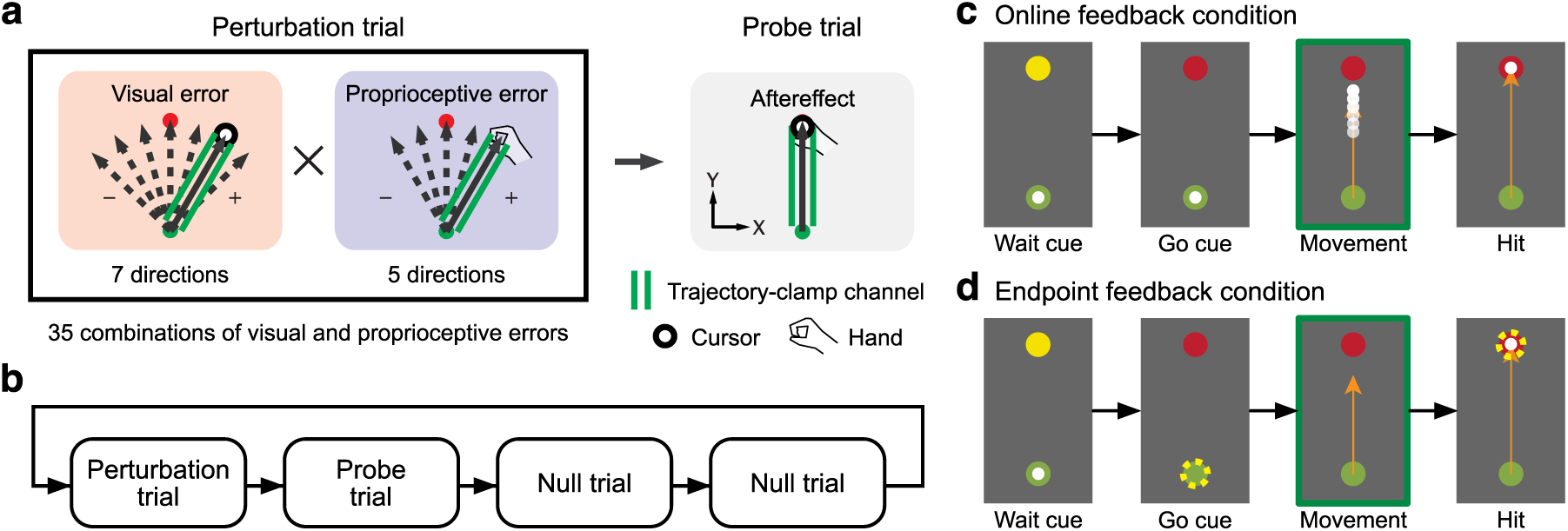
Procedures of Experiments 1 and 2. (**a**) Participants made reaching movements towards a front target (distance = 10 cm). In the perturbation trial, the force channel was used to provide a visual and/or proprioceptive perturbation. There were 35 combinations of perturbations in total (seven visual perturbations from −45° to 45° and five proprioceptive perturbations from −30° to 30°). In the subsequent probe trial, the force channel was used to measure the aftereffect. (**b**) The perturbation and probe trials were followed by two ordinary null trials to washout the effect of adaptation. This small block was repeated. (**c**) In Experiment 1 (Online feedback condition), the cursor was continuously visible during the reaching movement. (**d**) In Experiment 2 (Endpoint feedback condition), the cursor disappeared during the movement and reappeared only after completion of the movement.

### Aftereffects to each of visual and proprioceptive error

Fig. 2 illustrates the evolvement of lateral force with the movement time in the probe trials after only visual (Fig. 2a) or proprioceptive perturbation (Fig. 2b) was imposed. The presence of an aftereffect was confirmed as the lateral force in the direction opposite to the error imposed in the preceding perturbation trial. The aftereffect was quantified as the integrated lateral force over the time interval from the force onset to the time at the peak hand velocity (i.e., feedforward component: inset in Fig. 2c). Notably, not only the visual (Fig. 2c, one-way repeated measures ANOVA, F_(6,54)_ = 21.979, p = 6.101 × 10^−13^), but also the proprioceptive perturbation (Fig. 2d, F_(4,36)_ = 18.861, p = 1.934 × 10^-8^) can elicit aftereffects, although the cursor safely reached the target, indicating that the proprioceptive sensory prediction error was also used for motor adaptation^5,9^. Furthermore, as reported in previous studies^17–19,27^, the aftereffect did not increase linearly with the size of perturbation.

**Fig. 2.**
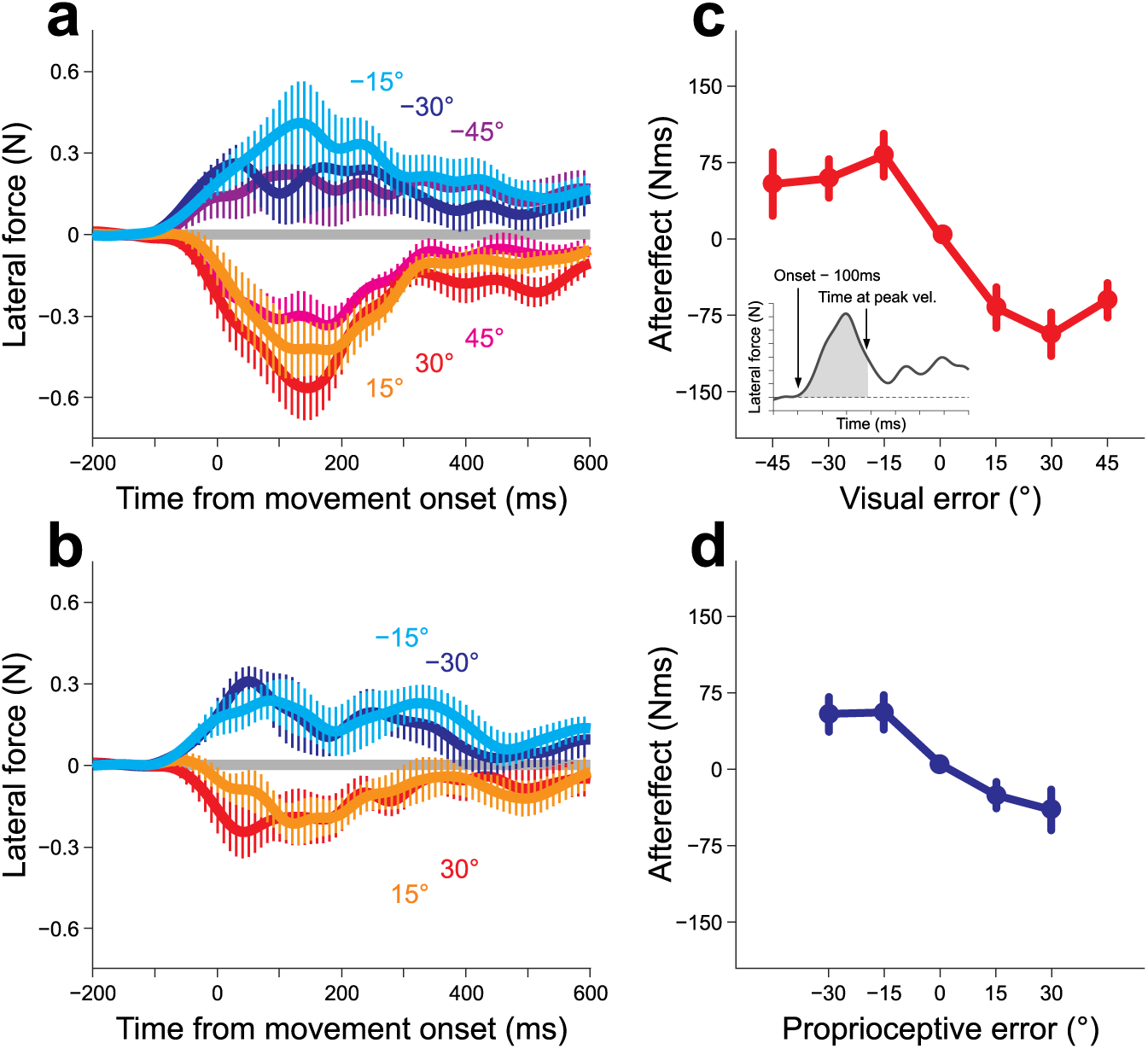
Aftereffects of visual or proprioceptive perturbation in Experiment 1. Time series of the lateral force exerted against the force channel in the probe trial after either visual (**a**) or proprioceptive perturbation (**b**) was imposed. The relationship between the aftereffect and the size of the visual (**c**) or proprioceptive (**d**) error. The aftereffects were quantified by the integrated lateral force from the force onset to the time of peak velocity (inset of **c**). The error bars represent the standard error across participants.

### Aftereffects of combinations of visual and proprioceptive errors

Fig. 3a-e illustrates the modulation of the lateral force induced by visual perturbation with additional proprioceptive perturbation. As indicated in Fig.3c (or Fig. 2a), the positive and negative visual perturbations induced negative and positive lateral force (aftereffects), respectively. However, when +30° proprioceptive perturbation was additionally imposed (Fig. 3a), the aftereffect curves for the negative visual perturbations (−15°, −30°, −45°) were considerably suppressed, while the aftereffect curves for the positive visual perturbation (15°, 30°, 45°) were only slightly altered. The suppression of the lateral force for the opposite visual and proprioceptive perturbation directions and relatively unchanged lateral force for the same perturbation directions was observed when the other amount of proprioceptive perturbations (−30°, −15°, and 15°) were additionally imposed (Fig. 3a-e).

**Fig. 3.**
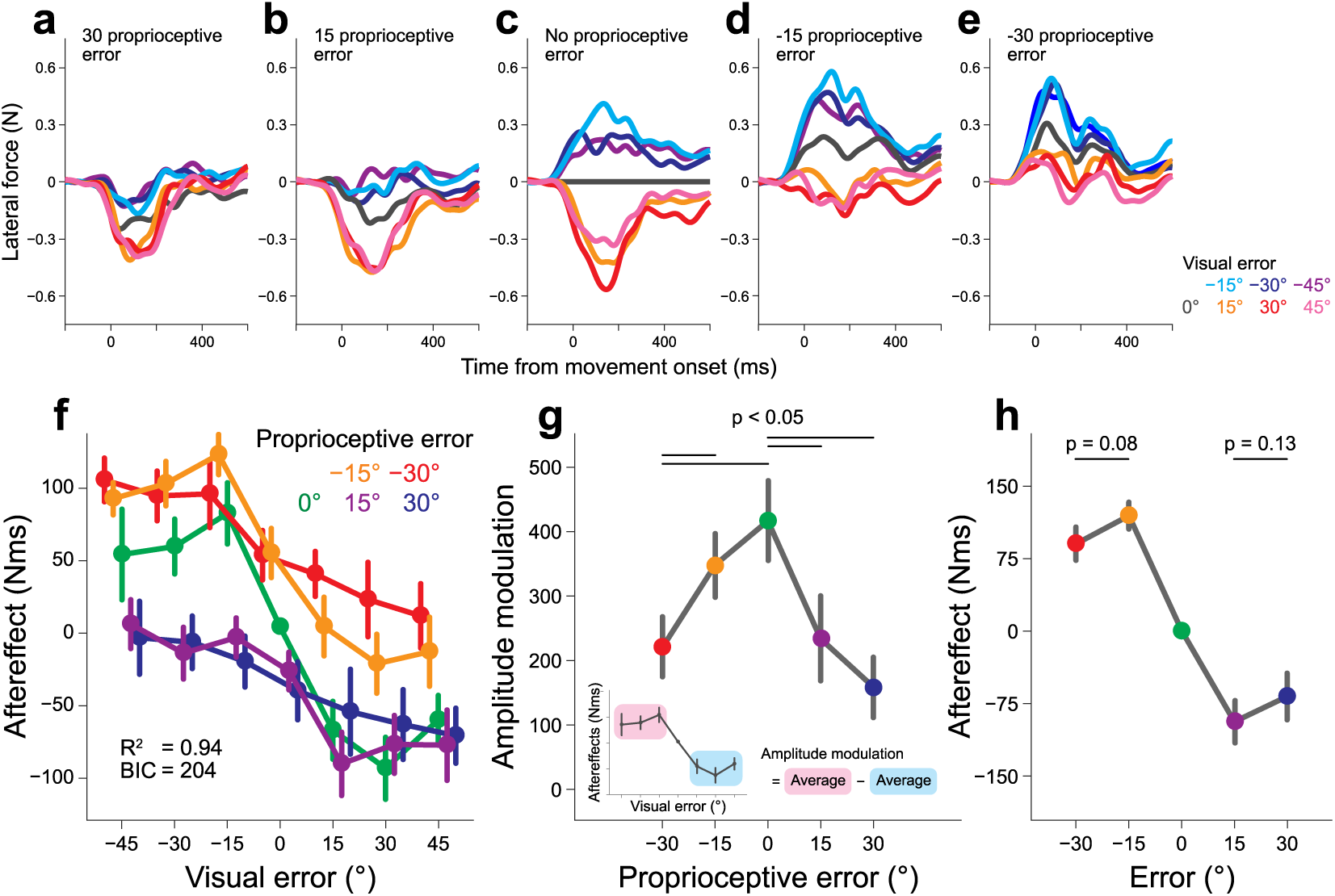
Aftereffects in Experiment 1 when various combinations of visual and proprioceptive perturbations were applied. Each panel presents the lateral force exerted against the force channel in the probe trials after seven different visual perturbations were imposed (proprioceptive perturbation = 30° [**a**], 15° [**b**], 0° [**c**], −15° [**d**], −30° [**e**]). (**f**) The dependence of the aftereffect on the size of the visual error. Each line represents the aftereffect when different proprioceptive errors are simultaneously imposed. (**g**) The effect of the size of the proprioceptive error on the amplitude of the aftereffect modulation of the visual error size. The amplitude was calculated as the difference between the aftereffects of positive and negative visual perturbations (inset). The presence of proprioceptive error decreased the amplitude of the modulation. The horizontal bars indicate the significant differences. (**h**) The aftereffect when the visual and proprioceptive errors were identical. Although the relevance of the error was maximal, saturation of the aftereffect with the error size was still observed. The error bars represent the standard error across participants.

Fig. 3f demonstrates the aftereffects of all 35 combinations of visual and proprioceptive perturbations. There was a significant interaction between visual and proprioceptive perturbations (two-way repeated measures ANOVA, F_(24,216)_ = 3.479, p = 5.233 × 10^−7^), indicating that the effect of proprioceptive perturbation on visual perturbation was not simply additive. Fig. 3f also suggests that the degree of modulation with visual perturbation size (i.e., the amplitude of each line) decreased with additional proprioceptive perturbations. To evaluate the size of the modulation, we calculated the difference between the aftereffects for positive visual perturbations (15°, 30°, and 45°) and those for negative visual perturbations (−15°, −30°, and −45°) (Fig. 3g). A one-way repeated measures ANOVA indicated that the size of the modulation significantly differed between the size of the proprioceptive perturbations (F_(4,36)_ = 10.27, p = 1.518 × 10^−7^). A post-hoc test revealed that the size of the modulation significantly decreased with the size of the additional proprioceptive perturbation (Fig. 3g; p < 0.05 by Bonferroni-Holm correction).

In the present experiment, there were five conditions in which the visual perturbation was identical to the proprioceptive perturbation (i.e., 0°, ±15°, ±30°). In these conditions, the relevance of the visual error was perfectly maintained, predicting that the aftereffects were not saturated with the size of the error^17^. However, as indicated in Fig. 3h, saturation of the aftereffects was still observed: the size of aftereffect for ±30° was not larger than that for ±15° (−30° vs −15°, t_(9)_ = 1.934, p = 0.0851; 30° vs 15°, t_(9)_ = 1.631, p = 0.1372, respectively).

### Possible integration mechanism: Divisive normalization

Previously proposed models were unlikely to explain our results (Fig. 3f-h) (see Supplementary Information). First, the highly complicated pattern of the aftereffects and the presence of a significant interaction between visual and proprioceptive factors (Fig. 3f) led us to reject the simple Bayesian integration model wherein the aftereffect is induced according to the error optimally integrating visual and proprioceptive errors^11,12^ and the aftereffect is a linear summation of those induced by visual and proprioceptive errors individually^19^. Second, the “relevance of the error”^17^ model was also rejected because the aftereffects did not linearly increase with the size of the error (Fig. 3h) even when the visual and proprioceptive errors were congruent (i.e., the relevance of the error was maximal).

We attempted to determine if the divisive normalization model (Eq. 1) could reasonably explain our results. Fig. 4a indicates *f*(*v*, *p*) obtained by fitting the model to the data using the least square method (R^2^ = 0.9423, *w*_*v*_ = 97.61, *w_p_* = 154.12, *k*_*v*_ = 1.99, *k_p_* = 7.73). Notably, this model captured the remarkable features of the experimental results. First, the size of the aftereffect did not linearly increase with the size of the error due to the presence of a normalization factor (Figs. 3f and 4a). Second, the size of the aftereffect modulation by the visual error decreased with the size of the proprioceptive error (Figs. 3g and 4b). Third, the relevance of the error was not related to the saturation of the aftereffect with the error size (Figs. 3h and 4c).

**Fig. 4.**
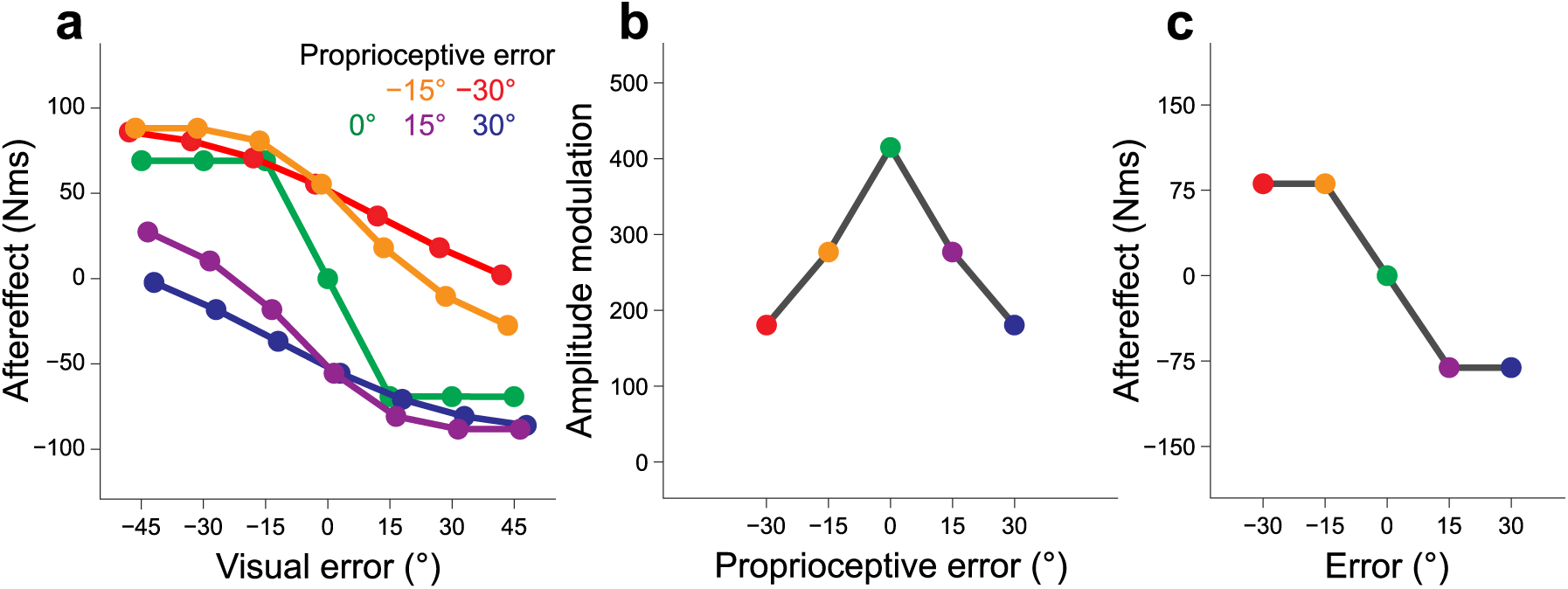
The divisive normalization model to explain the experimental results. (**a**) The divisive normalization model (Eq. 1) fit the experimental data and nicely reproduced the complicated pattern of the aftereffect of the visual and proprioceptive perturbations. The parameters were obtained using the least square method. The model also reproduced the reduction in the amplitude modulation with the proprioceptive perturbation (**b**) and the saturation of the aftereffect with the error size **(c)**.

### Effects of the presence and absence of online visual errors on the aftereffects

In Experiment 1, the participants were exposed to visual and proprioceptive errors during movement. Because multisensory integration can occur during movement^28–30^, the sensorimotor system can automatically generate online feedback responses to multisensory errors. Previous studies have suggested that the feedback responses function as a teaching signal for motor adaptation^31,32^. Thus, it is possible that the divisive normalization pattern in the aftereffects could reflect the feedback responses in the preceding perturbation trial. In order to examine this possibility, we quantified the feedback response as the integrated lateral force over the time interval from the time at the peak handle velocity to movement termination (i.e., feedback component: Fig. 5a-f). The degree of modulation of the feedback response with visual perturbation size seems relatively unchanged by the additional proprioceptive perturbation (Fig. 5f), which was contrasted with that of the aftereffect (Fig. 3f). This might indicate the smaller contribution of normalization factor in a divisive normalization model as reflected by smaller values of *k*_*v*_ and *k_p_* obtained by fitting the data with Eq. 1 (R^2^ = 0.9780, *w*_*v*_ = 3.20, *w_p_* = 24.66, *k*_*v*_ = 4.247 × 10^−15^, and *k_p_* = 9.068 × 10^−5^).

**Figure 5.**
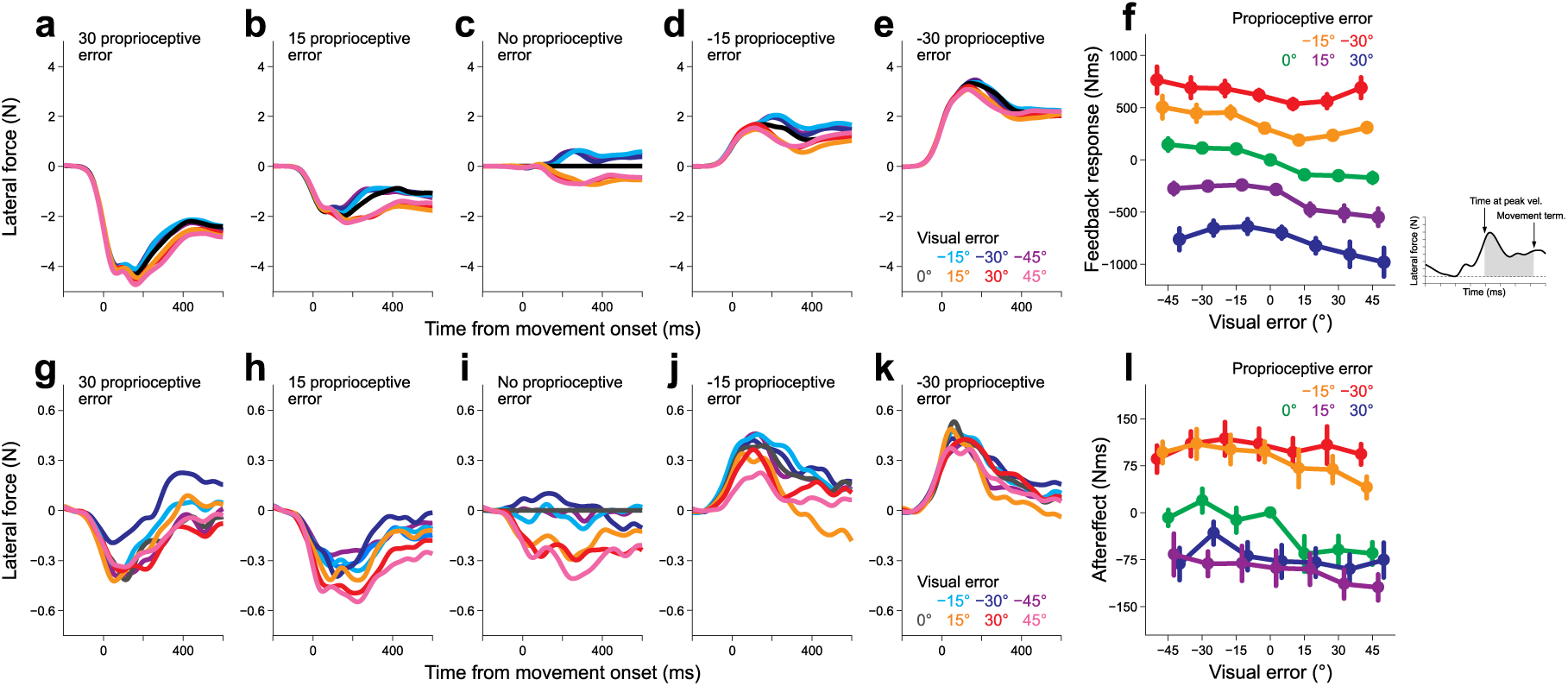
The online feedback response in Experiment 1 and the aftereffect in Experiment 2. The first five panels present the lateral force exerted against the force channel in the perturbation trial in which seven different visual perturbations were imposed (proprioceptive perturbation = 30° [**a**], 15° [**b**], 0° [**c**], −15° [**d**], −30° [**e**]). **(f)** The dependence of the feedback response on the size of the visual perturbation. Each line represents the aftereffect for different proprioceptive perturbations. The feedback response was quantified by the integrated force from the time of peak hand velocity to the movement offset (inset). The next five panels present the lateral force exerted against the force channel in the probe trials after seven different visual perturbations were imposed (proprioceptive perturbation = 30° [**g**], 15° [**h**], 0° [**i**], −15° [**j**], −30° [**k**]). (**l**) The dependence of the aftereffect on the size of the visual perturbation. Each line represents the aftereffect for different proprioceptive perturbations. The error bars represent the standard error across participants.

However, it should be noted that the feedback responses among the different proprioceptive perturbations could not be directly compared, because the movement directions were different. Thus, the difference in the pattern between the aftereffect (Fig. 3f) and the feedback response (Fig. 5f) cannot be solely ascribed to the involvement of different mechanisms. To further investigate this issue, we performed Experiment 2 in which the cursor was only visible immediately after movement completion (Fig. 1d). In this experiment, the visual information was not provided during reaching movements to eliminate the online feedback response to the visual perturbation. Nevertheless, the aftereffect still exhibited a divisive normalization pattern (R^2^ = 0.9403, *w*_*v*_ = 2.902, *w_p_* = 34.90, *k*_*v*_ = 0.0055, and *k_p_* = 0.1434, Fig. 5g-l). Therefore, the divisive normalization pattern of the aftereffects observed in Experiment 1 (Fig. 3f) could not be fully explained by the feedback response (Fig. 5f).

### The stage of integration of visual and proprioceptive information

A remaining question was at which stage the visual and proprioceptive information are integrated. There are two possibilities: the modality-shared motor memory model (Fig. 6a) or the modality-specific motor memory model (Fig. 6b). The modality-shared motor memory model (Fig. 6a) assumes that visual and proprioceptive sensory prediction errors are used to estimate a single integrated error according to a divisive normalization mechanism and this integrated error is used for updating a motor memory. In contrast to this ordinary interpretation of multisensory integration, the modality-specific motor memory model (Fig. 6b) assumes that each modality has its own motor memory separately updated, and then the outputs from these motor memories are integrated (See also Supplementary Information).

**Fig. 6.**
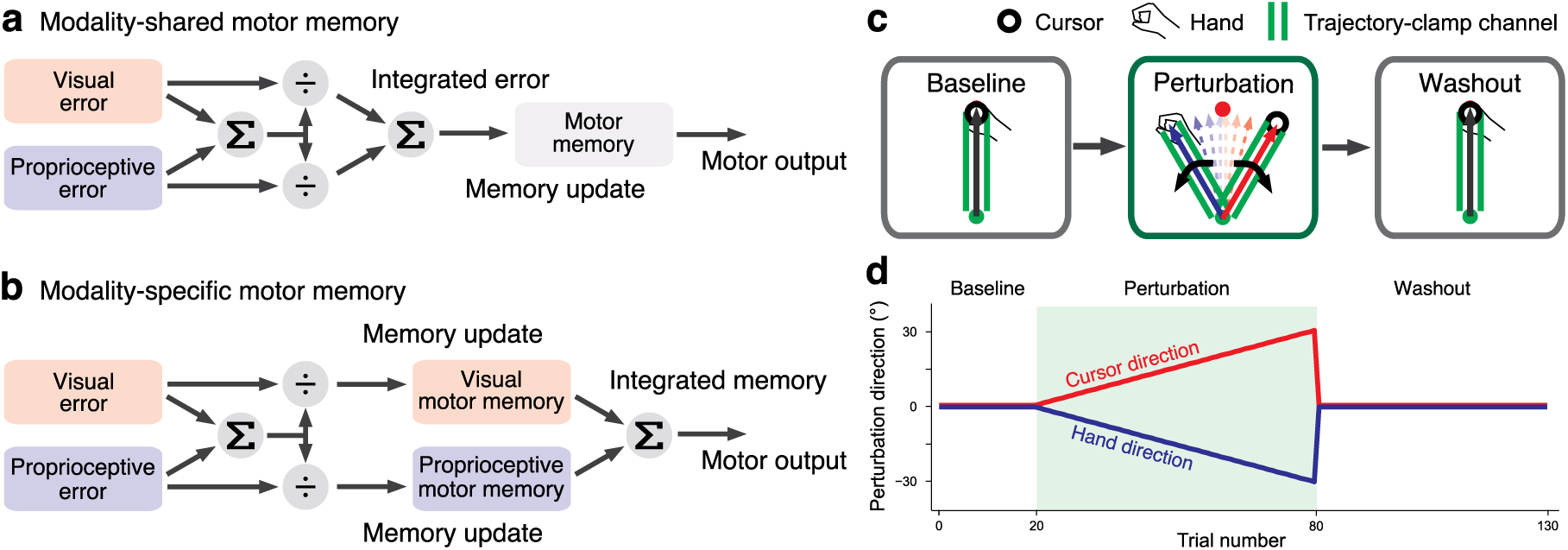
Possible integration scheme and the procedure of Experiment 3. In the modality-shared motor memory model (**a**), visual and proprioceptive errors are integrated by the divisive normalization mechanism to obtain the integrated error information. Motor memory is updated by the integrated error. In contrast, in the modality-specific motor memory model (**b**), visual and proprioceptive memories are independently created. **(c, d)** Experiment 3 was designed to determine which integration scheme was more likely. After 20 force channel trials to the forward target, gradually increasing visual and proprioceptive perturbations were imposed in the opposite direction (60 trials). This perturbation phase was followed by a washout phase in which the force channel trials to the forward target were repeated (50 trials).

Experiment 3 was designed to determine which model was more likely (Fig. 6c, d). Fourteen participants (12 male and 2 female, 20-25 years old) reached towards a frontal target while receiving the gradually increasing visual and proprioceptive perturbations in the opposite directions for 60 trials (perturbation phase). We measured the aftereffects during the following 50 force channel trials toward the same front target (washout phase: both visual and proprioceptive perturbations were turned off) by quantifying the lateral force against the force channel. The participants were instructed to aim towards the frontal target consistently throughout the experiment.

The modality-shared motor memory model predicts that the aftereffect after the completion of perturbation phase (i.e., the memory content), though it is expected to be suppressed due to the perturbations in the opposite directions, should merely decay during washout trials (Fig. 7a). In contrast, the modality-specific motor memory model could demonstrate a totally different behavior when a particular condition is met. Since this model has modality-specific motor memories, each motor memory could develop the memory content in the opposite directions (Fig. 7b). If the time-constant of trial-dependent decay is different between the memories, the motor memory of slower modality could emerge as the washout trials progress (Fig. 7b).

**Fig. 7.**
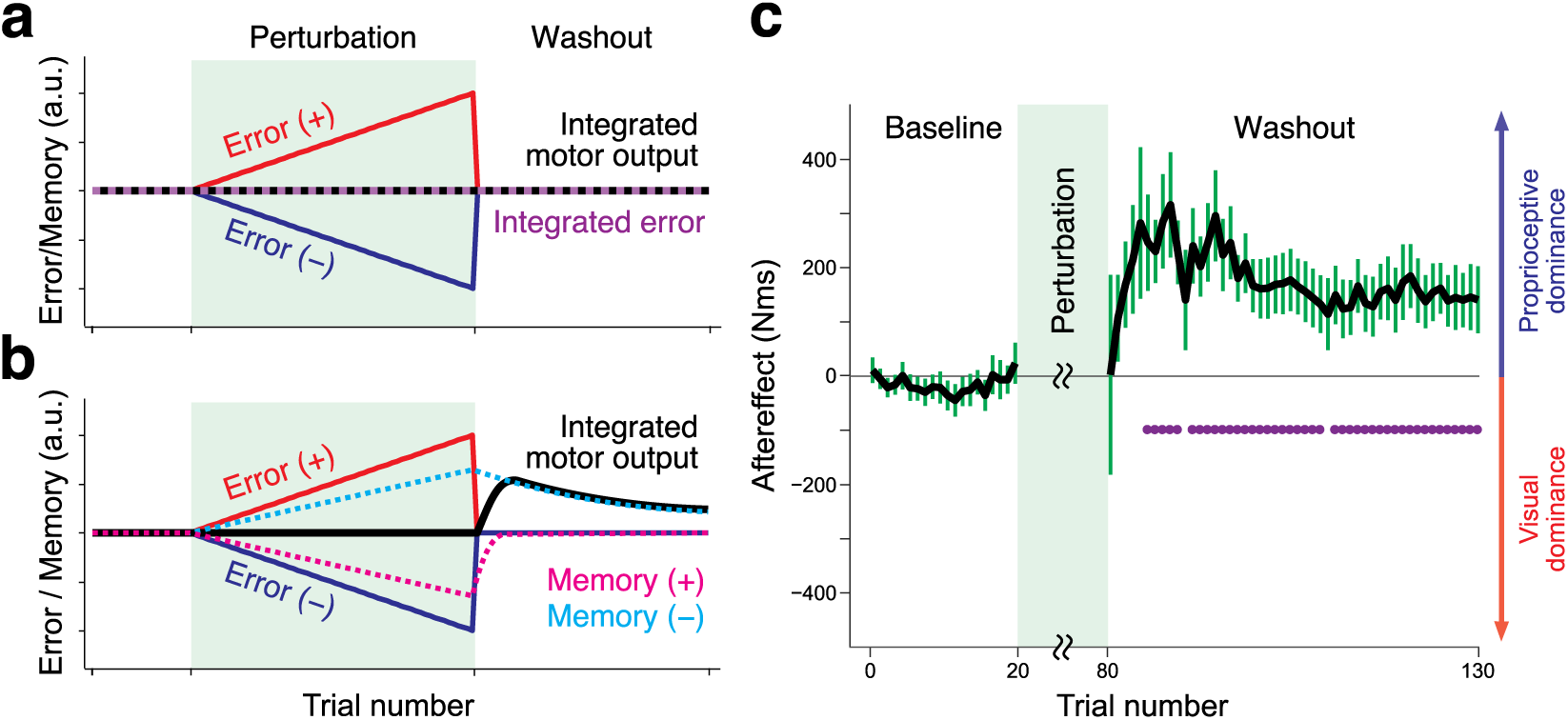
Model predictions and experimental data of Experiment 3. The prediction of lateral force during the washout phase by the modality-shared motor memory model **(a)** and by the modality specific motor memory model **(b)**. **(c)** The experimental results indicated that the aftereffect at the beginning of the washout phase was indistinguishable from that of the baseline trials. However, as the trials progressed, an aftereffect that was significantly greater than zero appeared and then decayed. The error bars represent the standard error across participants. The purple dots indicate the trials in which aftereffects that were significantly greater than zero (as determined by a t-test with false discovery correction) were observed.

As expected from the opposite directions of the visual and proprioceptive perturbations, significant aftereffects were absent in the beginning of the washout trials (Fig. 7c: t_(13)_ = 0.013, p = 0.9898). However, as the washout phase progressed, the aftereffect began to emerge in the direction opposite to that of the proprioceptive perturbation (i.e., in the direction of the visual perturbation), reached a maximum, and then decayed (Fig. 7c). Therefore, the experimental result supported the modality-specific motor memory model (Figs. 6b and 7b). A similar trial-dependent aftereffect pattern could be reproduced by the state space model implementing the memory integration mechanism (Supplementary Information).

## DISCUSSION

We investigated the utilization of visual and proprioceptive information in motor adaptation by precisely imposing various perturbation combinations using a force channel method. The present results indicate that the aftereffect was dependent on the visual and proprioceptive errors in a highly complicated manner (Figs. 2 and 3), and this pattern was reasonably explained by the divisive normalization mechanism (Fig. 4). Furthermore, we demonstrated that the visual and proprioceptive motor memories were likely created independently (Figs. 6 and 7).

### Comparison with previously proposed mechanisms of visual and proprioceptive information integration

A previous study suggested that visual and proprioceptive information are independently processed for motor adaptation^19^. According to this hypothesis, the aftereffect should be a summation of the aftereffects induced individually by visual or proprioceptive perturbation. However, this was not the case in the current study, as clearly demonstrated by the presence of a significant interaction in the aftereffects between the visual and proprioceptive perturbations (Fig. 3f). The contradiction was likely derived from the fact that our experiment imposed a wider range of visual and proprioceptive errors; in the previous study, the error size was < 15° (estimated from their data) and the combinations of opposite perturbation directions were not tested.

Another important theory of multisensory integration is the Bayesian theory. According to this theory, the sensory prediction error can be estimated by a linear summation of the visual and proprioceptive errors and the relative weights are determined by the signal reliabilities. More reliable sensory information has a greater weight and the estimated error is biased toward more reliable sensory information^10–12,33^. Clearly, the simple Bayesian model cannot explain the saturation of aftereffects with error size (Supplementary Information).

One plausible solution is to assume that the reliability of sensory information depends on the signal intensity^34,35^. Consider the case in which the size of the visual perturbation increases while the proprioceptive perturbation is maintained at zero. If the reliability of the visual error information decreases with the error size, the integrated error should be more biased toward the hand direction (i.e., proprioception). Thus, the aftereffect, which reflects the estimated error size, should not increase with the visual perturbation size, which explains the saturation of aftereffect with perturbation size (Fig. 2a, b).

However, the model’s prediction was not consistent with our results in the several points (Supplementary Information). First, when the sizes of visual and proprioceptive perturbation are identical, the bias of the estimated error toward a more reliable error was absent, which should eliminate the saturation effect. However, reduction in the aftereffect with error size was still observable^19^ (Fig. 3h). Second, the modulation of the aftereffect with the visual perturbation size decreased when proprioceptive perturbation was additionally introduced (Fig. 3g). However, this modified Bayesian theory provides the opposite prediction that the modulation increased. Taken together, the mechanism based on the Bayesian theory cannot explain our results.

### The aftereffect pattern is explained by a divisive normalization mechanism

Divisive normalization is proposed as a canonical neural computation mechanism in the brain^20^. The explanatory power extends from the neural activities in the retina, primary visual cortex, primary auditory cortex, olfactory system, middle temporal area, and parietal cortex, to the psychophysical features of perceptual decision making, value-based decision making, face-attractiveness evaluation, and attention^21,22,36–44^. Many previous studies have also incorporated this mechanism into the neural network model^45–48^.

We found that the divisive normalization model could reasonably reproduce the complicated dependence of the aftereffect on visual and proprioceptive perturbations. Ohshiro et al.^21,22^ demonstrated that the divisive normalization mechanism can account for the response of multisensory neurons encoding vestibular and visual signals. The neural response to both modalities signals is smaller than the summation of the activity in response to sensory signals presented individually (cross-modal suppression), which is considered to be the signature of the divisive normalization mechanism. In our experiment, the aftereffect demonstrated a cross-modal suppression-like phenomenon; the aftereffect of combinations of visual and proprioceptive perturbations was smaller than the summation of the aftereffects of each perturbation (e.g., | *f* (30,30)| < |*f*(30, 0)| + |*f*(0, 30)| in Fig. 3f).

We believe that this is the first behavioral demonstration of the divisive normalization mechanism accounting for the motor adaptation to visual and proprioceptive errors. This mechanism can explain not only why the aftereffects saturate with error size, but also the complicated dependence on visual and proprioceptive errors (Fig. 3f and Fig. 4a). However, there are several remaining issues that need to be investigated. First, the link between the response at the behavior level and the response at the neuronal circuit level remains unknown. A recent study^49^ has reported that the divisive normalization mechanism accounts for the neuronal response in the superior parietal lobule, which receives visual and proprioceptive signals, while monkeys performed reaching movement. Although the post parietal area is known to be involved in visuomotor control and learning^50–53^, it is still difficult to comprehend how this knowledge can be unified with our finding.

Second, although the functional significance has been proposed from the perspective of efficient coding of sensory information^20,54^, the functional role in the motor adaptation system is unclear. Shadmehr et al. demonstrated that adaptation to a force field in people with autism predominantly relied on proprioception^55,56^. They also illustrated that aftereffects to combinations of them showed a complicated modulation between healthy and autism people although aftereffects specific to visual or proprioceptive perturbation were likely to be parallelly shifted^57^, implying the possibility that the multisensory integration pattern by divisive normalization is peculiar in people with autism. Indeed, a recent study proposed that disorders in people with autism are related to the alteration of the divisive normalization pattern derived from the imbalance between excitation and inhibition in neuronal circuits^58^. Studies investigating the peculiar characteristics of motor adaptation in patients with autism or neurological disorders might be beneficial to our understanding of the functional significance of divisive normalization in the motor adaptations and the neuronal architecture implementing divisive normalization.

### Influence of online and endpoint feedback

According to the feedback error learning^31^, the feedback motor command is used as a teaching signal to modify the motor command in the subsequent trial^32^. Thus, it is possible that the divisive normalization pattern in the aftereffect reflects the feedback response pattern. However, the patterns were considerably different between the aftereffect (Fig. 3f) and the feedback response (Fig. 5f). Furthermore, the aftereffect in the endpoint feedback condition exhibited the divisive normalization pattern (Fig. 5l). Therefore, the divisive normalization pattern in the aftereffect cannot be ascribed either to the feedback response or to the endpoint error processing.

Notably, the considerably different patterns between Figs. 3f and 5l implied that the neuronal processing of online and endpoint errors for motor adaptation are somehow dissociated^59,60^. The sensitivity of the aftereffect to visual error was considerably suppressed in the endpoint error feedback condition (Fig. 3f vs Fig. 5l), indicating that the influence of the online proprioceptive error was dominant in the endpoint feedback condition and that the contribution of online visual feedback was greater than that of endpoint visual feedback in the production of the divisive normalization pattern. Further studies are necessary to clarify how these two types of feedback are cooperatively involved in creating the divisive normalization pattern in the aftereffect.

### Modality-specific motor memory

Previous studies have investigated whether adaptation of visual rotation (kinematic) and of a novel force field (dynamic) is dependently^61^ or independently accomplished^62^. The results of Experiment 3 are relevant to this problem because the findings illustrated that motor memories for vision and proprioception are created independently, and integrated at the output stages (Fig. 6b). The notion of separate motor memories for visual and proprioception has been previously proposed^26^, but their conclusion that visual motor memory was dominant in motor adaptation is not consistent with our finding that proprioceptive error had a substantial influence on the aftereffect of visual error. This contradiction might result from their assumption that motor memories are updated by the linear summation of both modality errors, which differed from our scheme that indicated nonlinear integration.

The emergence of the aftereffect during the washout phase likely indicated that the memory for proprioception has a slower time constant than does the memory for vision (Fig. 7). This might be consistent with the previous observation that, even after adaptation to visual rotation, the elimination of visual feedback led the hand to move to the preadapted direction^63^, implying that the adaptation of proprioception is slower. It has been recognized that the adaptation to a novel dynamical environment is accomplished by slow and fast adaptation processes^64^. Implicit and explicit motor learning have been reported to correspond to slow and fast processes, respectively^65^, but our results indicate a different possibility that slow and fast processes are related to proprioception and vision, respectively.

In conclusion, we demonstrated that the motor adaptation system integrates visual and proprioceptive errors information by a divisive normalization mechanism. Furthermore, we found evidence that the motor memory for each sensory modality was formed separately and the outputs from these memories were integrated. These results provide a novel view of the utilization of different sensory modality signals in motor control and adaptation.

## METHODS

### Ethics statement

Thirty-four right-handed participants (26 males and eight females, aged 20-31 years) volunteered to participate in the three experiments. Before the experiments started, we fully explained the experimental procedures and written informed consent was obtained from all participants. The ethics committee of the University of Tokyo approved the experiments.

### General settings

The participants performed reaching movements while holding a manipulandum (KINARM End-point Lab, BKIN Technologies, Kingston, ON, Canada). To reduce unwanted wrist movement and postural fluctuation, their arms were constrained by a brace and supported in a horizontal plane with a spring sling. The green target (diameter: 10 mm) was located 10 cm from the start position immediately in front of the participant. After 0.5 – 0.7 seconds, the target color turned to magenta, which was the “go” cue. The participants were asked to move the handle of the KINARM robot straight towards the target as smoothly as possible. In Experiments 1 and 3, the cursor (diameter: 10 mm) representing the position of the handle was continuously visible, while the cursor was visible only after the reaching movement was complete in Experiment 2. At the end of each trial, a warning message (“fast” or “slow”) appeared when the movement speeds were too fast (> 450 mm/s) or too slow (< 250 mm/s). The participants maintained the hand position at the end of the movement until the robot automatically returned the handle to the starting position (for 1.5 s). The force channel method in the perturbation trial (see the next section) could not allow the participants to correct the movement trajectories. Thus, the cursor could never reach the target under the presence of visual perturbations in Experiments 1 and 3. Similarly, in Experiment 2, the participants were unable to know the movement distance until completion of the movement. The participants practiced so that they could terminate the movement at the appropriate distance (the actual movement distance was 9.85 ± 0.1 cm).

### Experiments 1 and 2

In Experiment 1 (ten participants: eight males and two females, aged 21-25 years) and Experiment 2 (ten participants: six males and four females, aged 21-31 years), we examined a single-trial motor adaptation induced by 35 combinations of seven visual (± 45°, ± 30°, ± 15°, and 0°; Fig. 1a red) and five proprioceptive perturbations (± 30°, ± 15°, and 0°; Fig. 1a blue). These perturbations were applied by constraining the hand trajectory in a straight line using the force channel method^25^. The force channel was created by a virtual spring (6000 N/m) and dumper (100 N/[m/s]) in the perpendicular direction to the straight path of the movement. This procedure enabled us to completely control the sizes of the visual and proprioceptive error in the same angular unit (°). In the following probe trial, the lateral force against the force channel was measured to evaluate the aftereffect (Fig. 1a gray). One set consisted of a perturbation trial (one of 35 combinations was pseudo-randomly selected) and a probe trial followed by two ordinary null trials (without the force channel) to washout the adaptation effect (Fig. 1b). The cursor was always visible (online feedback: Fig. 1c) in Experiment 1, while the cursor was only visible after the completion of the reaching movement (endpoint feedback: Fig. 1d) in Experiment 2.

### Experiment 3

We then aimed to determine the stage at which the visual and proprioceptive information is integrated. There are two potential models: the modality-shared motor memory model (Fig. 6a) or the modality-specific motor memory model (Fig. 6b). The modality-shared motor memory model (Fig. 6a) assumes that visual and proprioceptive sensory prediction errors are used to estimate a single integrated error according to a divisive normalization mechanism, and this integrated error is used for updating a motor memory. In contrast to this ordinary interpretation of multisensory integration, the modality-specific motor memory model (Fig. 6b) assumes that the motor memories of each modality are separately updated, and then the outputs from these motor memories are integrated (see Supplementary Information).

Experiment 3 was designed to determine which model was more likely. Fourteen participants (12 males and 2 females, aged 20-25 years) performed 131 reaching movements in Experiment 3 (Fig. 6c, d). After being familiarized to the procedures, they performed reaching movements with the virtual channel in the baseline session (20 trials). The participants were then exposed to the visual and proprioceptive perturbations simultaneously in the perturbation session (61 trials). The degrees of perturbation were gradually increased at a rate of 0.5° to 30°. The directions of the perturbations were opposite and counter-balanced across the participants. In the subsequent washout session, they again performed reaching movements with the virtual channel (50 trials).

The modality-shared motor memory model predicts that the aftereffect after the completion of the perturbation phase (i.e., the memory content), although it is expected to be suppressed due to perturbations in the opposite directions, should merely decay during washout trials (Fig. 7a). In contrast, the modality-specific motor memory model could predict completely different behavior when a particular condition is met. Since this model involves modality-specific motor memories, each motor memory could develop the memory content in the opposite directions (Fig. 7b). If the time-constant of trial-dependent decay differs between the memories, the motor memory of the slower modality could emerge as the washout trials progress (Fig. 7b).

### Data analysis

The position and force data of the handle were sampled at a rate of 1000 Hz and filtered by a 4th ordered zero-lag Butterworth filter with cutoff a frequency of 10 Hz. The position of the handle was numerically differentiated to obtain the handle velocity. We quantified the aftereffect in the probe trials as the integrated lateral force over the time interval from the force onset to the time at the peak handle velocity. We also quantified the amount of online feedback force in the perturbation trials as the integrated lateral force over the time interval from the time at the peak handle velocity to movement termination.

## Acknowledgement

We thank Professor Ken Takiyama and members of the Nozaki laboratory for their helpful comments and suggestions, and Asako Munakata for coordinating experiments. This study was supported by a grant from the Japan Society for the Promotion of Science Research Fellowships for Young Scientists to TH (17J02601) by a KAKENHI (17H00874) to DN.

## References

1. Shadmehr, R., Smith, M. A. & Krakauer, J. W. Error Correction, Sensory Prediction, and Adaptation in Motor Control. Annu. Rev. Neurosci. 33, 89–108 (2010).

2. Krakauer, J. W., Pine, Z., Ghilardi, M. & Ghez, C. Learning of visuomotor transformations for vectorial planning of reaching trajectories. J. Neurosci. 20, 8916–24 (2000).

3. Shadmehr, R. & Mussa-Ivaldi, F. A. Adaptive representation of dynamics during learning of a motor task. J. Neurosci. 14, 3208–3224 (1994).

4. Lackner, J. R. & Dizio, P. Rapid adaptation to Coriolis force perturbations of arm trajectory. J. Neurophysiol. 72, 299–313 (1994).

5. Franklin, D. W., So, U., Burdet, E. & Kawato, M. Visual Feedback Is Not Necessary for the Learning of Novel Dynamics. PLoS ONE 2, e1336 (2007).

6. Dizio, P. & Lackner, J. R. Congenitally blind individuals rapidly adapt to coriolis force perturbations of their reaching movements. J. Neurophysiol. 84, 2175–80 (2000).

7. Bernier, P., Chua, R., Bard, C. & Franks. I. M. Updating of an internal model without proprioception: a deafferentation study. NeuroReport 18, 1421–5 (2006).

8. Yousif, N., Cole, J., Rothwell, J. & Diedrichsen J. Proprioception in motor learning: lessons from a deafferented subject. Exp. Brain Res. 233, 2449–59 (2015).

9. Pipereit, K., Bock, O. & Vercher J. L. The contribution of proprioceptive feedback to sensorimotor adaptation. Exp. Brain Res. 174, 45–52 (2006).

10. Ernst, M. O. & Banks, M. S. Humans integrate visual and haptic information in a statistically optimal fashion. Nature 415, 429 (2002).

11. van Beers, R. J., Sittig, A. C. & Gon, J. J. Integration of proprioceptive and visual position-information: An experimentally supported model. J. Neurophysiol. 81, 1355–64 (1999).

12. van Beers, R. J., Wolpert, D. M. & Haggard, P. When Feeling Is More Important Than Seeing in Sensorimotor Adaptation. Curr. Biol. 12, 834–837 (2002).

13. Kagerer, F. A., Contreras-Vidal, J. L. & Stelmach, G. E. Adaptation to gradual as compared with sudden visuo-motor distortions. Exp. Brain Res. 115, 557–61 (1997).

14. Mazzoni, P. & Krakauer, J. W. An Implicit Plan Overrides an Explicit Strategy during Visuomotor Adaptation. J. Neurosci. 26, 3642–3645 (2006).

15. Hirashima, M. & Nozaki, D. Distinct motor plans form and retrieve distinct motor memories for physically identical movements. Curr. Biol. 22, 432–6 (2012).

16. Hayashi, T., Yokoi, A., Hirashima, M. & Nozaki, D. Visuomotor Map Determines How Visually Guided Reaching Movements are Corrected Within and Across Trials. eNeuro 3, (2016).

17. Wei, K. & Körding, K. Relevance of Error: What Drives Motor Adaptation? J. Neurophysiol. 101, 655–664 (2009).

18. Kasuga, S., Hirashima, M. & Nozaki, D. Simultaneous Processing of Information on Multiple Errors in Visuomotor Learning. PLoS ONE 8, e72741 (2013).

19. Marko, M. K., Haith, A. M., Harran, M. D. & Shadmehr, R. Sensitivity to prediction error in reach adaptation. J. Neurophysiol. 108, 1752–63 (2012).

20. Carandini, M. & Heeger, D. J. Normalization as a canonical neural computation. Nat. Rev. Neurosci. 13, 51–62 (2011).

21. Ohshiro, T., Angelaki, D. E. & DeAngelis, G. C. A normalization model of multisensory integration. Nat. Neurosci. 14, 775–82 (2011).

22. Ohshiro, T., Angelaki, D. E. & DeAngelis, G. C. A Neural Signature of Divisive Normalization at the Level of Multisensory Integration in Primate Cortex. Neuron 95, 399–411 (2017).

23. Webb, R., Glimcher, P. W. & Louie, K. Rationalizing Context-Dependent Preferences: Divisive Normalization and Neurobiological Constraints on Choice. SSRN Electron. J. (2014).

24. Ren, M., Liao, R., Urtasun, R., Sinz, F. H. & Zemel, R. S. Normalizing the Normalizers: Comparing and Extending Network Normalization Schemes. arXiv (2016).

25. Scheidt, R. A., Reinkensmeyer, D. J., Conditt, M. A., Rymer, W. Z. & Mussa-Ivaldi, F. A. Persistence of motor adaptation during constrained, multi-joint, arm movements. J. Neurophysiol. 84, 853–62 (2000).

26. Judkins, T. & Scheidt, R. A. Visuo-proprioceptive interactions during adaptation of the human reach. J. Neurophysiol. 111, 868–87 (2014).

27. Körding, K. P. & Wolpert, D. M. The loss function of sensorimotor learning. Proc. Natl. Acad. Sci. USA 101, 9839–42 (2004).

28. Scott, S. H. A Functional Taxonomy of Bottom-Up Sensory Feedback Processing for Motor Actions. Trends Neurosci. 39, 512–26 (2016).

29. Crevecoeur, F., Munoz, D. P. & Scott, S. H. Dynamic Multisensory Integration: Somatosensory Speed Trumps Visual Accuracy during Feedback Control. J. Neurosci. 36, 8598–611 (2016).

30. Oostwoud Wijdenes, L. & Medendorp, W. P. State Estimation for Early Feedback Responses in Reaching: Intramodal or Multimodal? Front. Integr. Neurosci. 11, 38 (2017).

31. Kawato, M., Furukawa, K. & Suzuki, R. A hierarchical neural-network model for control and learning of voluntary movement. Biol. Cybern. 57, 169–185 (1987).

32. Albert, S. T. & Shadmehr, R. The Neural Feedback Response to Error As a Teaching Signal for the Motor Learning System. J. Neurosci. 36, 4832–4845 (2016).

33. Körding, K. P. et al. Causal Inference in Multisensory Perception. PLoS ONE 2, e943 (2007).

34. Jones, K. E., Hamilton, A. F. & Wolpert, D. M. Sources of signal-dependent noise during isometric force production. J. Neurophysiol. 88, 1533–44 (2002).

35. Goris, R. L., Movshon, A. J. & Simoncelli, E. P. Partitioning neuronal variability. Nat. Neurosci. 17, 858–65 (2014).

36. Heeger, D. J. Normalization of cell responses in cat striate cortex. Vis. Neurosci. 9, 181–97 (1992).

37. Carandini, M., Heeger, D. J. & Movshon, J. A. Linearity and normalization in simple cells of the macaque primary visual cortex. J. Neurosci. 17, 8621–44 (1997).

38. Simoncelli, E. P. & Heeger, D. J. A model of neuronal responses in visual area MT. Vision Res. 38, 743–61 (1998).

39. Busse, L., Wade, A. R. & Carandini, M. Representation of concurrent stimuli by population activity in visual cortex. Neuron 64, 931–42 (2009).

40. Reynolds, J. H. & Heeger, D. J. The Normalization Model of Attention. Neuron 61, 168–85 (2009).

41. Olsen, S. R., Bhandawat, V. & Wilson, R. I. Divisive normalization in olfactory population codes. Neuron 66, 287–99 (2010).

42. Louie, K., Grattan, L. E. & Glimcher, P. W. Reward Value-Based Gain Control: Divisive Normalization in Parietal Cortex. J. Neurosci. 31, 10627–10639 (2011).

43. . Louie, K., Khaw, M. W. & Glimcher, P. W. Normalization is a general neural mechanism for context-dependent decision making. Proc. Natl. Acad. Sci. USA 110, 6139–6144 (2013).

44. Furl, N. Facial-Attractiveness Choices Are Predicted by Divisive Normalization. Psychol. Sci. 27, 1379–1387 (2016).

45. Deneve, S., Latham, P. E. & Pouget, A. Reading population codes: a neural implementation of ideal observers. Nat. Neurosci. 2, 740–5 (1999).

46. Deneve, S., Latham, P. E. & Pouget, A. Efficient computation and cue integration with noisy population codes. Nat. Neurosci. 4, 826–831 (2001).

47. Verstynen, T. & Sabes, P. N. How each movement changes the next: an experimental and theoretical study of fast adaptive priors in reaching. J. Neurosci. 31, 10050–9 (2011).

48. Orhan, A. E. & Ma, W. J. Efficient probabilistic inference in generic neural networks trained with non-probabilistic feedback. Nat. Commun. 8, 138 (2017).

49. Shi, Y., Apker, G. & Buneo, C. A. Multimodal representation of limb endpoint position in the posterior parietal cortex. J. Neurophysiol. 109, 2097–107 (2013).

50. Tanaka, H., Sejnowski, T. J. & Krakauer, J. W. Adaptation to visuomotor rotation through interaction between posterior parietal and motor cortical areas. J. Neurophysiol. 102, 2921–32 (2009).

51. Mutha, P. K., Sainburg, R. L. & Haaland, K. Y. Critical neural substrates for correcting unexpected trajectory errors and learning from them. Brain 134, 3647–61 (2011).

52. Bédard, P. & Sanes, J. N. Brain representations for acquiring and recalling visual-motor adaptations. NeuroImage 101, 225–35 (2014).

53. Haar, S., Donchin, O. & Dinstein, I. Dissociating Visual and Motor Directional Selectivity Using Visuomotor Adaptation. J. Neurosci. 35, 6813–6821 (2015).

54. Beck, J. M., Latham, P. E. & Pouget, A. Marginalization in neural circuits with divisive normalization. J. Neurosci. 31, 15310–9 (2011).

55. Haswell, C. C., Izawa, J., Dowell, L. R., Mostofsky, S. H. & Shadmehr, R. Representation of internal models of action in the autistic brain. Nat. Neurosci. 12, 970–2 (2009).

56. Izawa, J. et al. Motor learning relies on integrated sensory inputs in ADHD, but over-selectively on proprioception in autism spectrum conditions. Autism Res. 5, 124–36 (2012).

57. Marko, M. K. et al. Behavioural and neural basis of anomalous motor learning in children with autism. Brain 138, 784–797 (2015).

58. Rosenberg, A., Patterson, J. S. & Angelaki, D. E. A computational perspective on autism. Proc. Natl. Acad. Sci. USA 112, 9158–65 (2015).

59. Saijo, N. & Gomi, H. Multiple Motor Learning Strategies in Visuomotor Rotation. PLoS ONE 5, e9399 (2010).

60. Izawa, J. & Shadmehr, R. Learning from Sensory and Reward Prediction Errors during Motor Adaptation. PLoS Comput. Biol. 7, e1002012 (2011).

61. Krakauer, J. W., Ghilardi, M.-F. & Ghez, C. Independent learning of internal models for kinematic and dynamic control of reaching. Nat. Neurosci. 2, 1026–31 (1999).

62. Tong, C., Wolpert, D. M. & Flanagan, R. J. Kinematics and dynamics are not represented independently in motor working memory: evidence from an interference study. J. Neurosci. 22, 1108–13 (2002).

63. Saijo, N. & Gomi, H. Effect of visuomotor-map uncertainty on visuomotor adaptation. J. Neurophysiol. 107, 1576–85 (2012).

64. Smith, M. A., Ghazizadeh, A. & Shadmehr, R. Interacting Adaptive Processes with Different Timescales Underlie Short-Term Motor Learning. PLoS Biol. 4, e179 (2006).

65. McDougle, S. M., Bond, K. M. & Taylor, J. A. Explicit and implicit processes constitute the fast and slow processes of sensorimotor learning. J. Neurosci 35, 9568–9579 (2015).

