## Supplementary Information for "Divisively normalized integration of multisensory error information develops motor memories specific to vision and proprioception"


### 1. Aftereffect pattern predicted by various computational models

The aftereffect obtained in our experiment (Fig. 3f-h) demonstrated the complicated interaction of various combinations of visual and proprioceptive errors. Fig. S1 summarizes the results of data fitting with possible candidate models (coefficient of determination and Bayesian Information Criterion (BIC)), indicating that the divisive normalization model is the most appropriate for explaining the complicated aftereffect pattern. Here, we describe the details of the models and explain why these models failed to reproduce the aftereffect pattern.

#### The optimal estimation of the error

Previous studies[1,2] have demonstrated that the hand location is optimally estimated using visual and proprioceptive information. If the same mechanism is used to obtain the sensory prediction error for motor adaptation, the integrated error can be obtained by a linear summation of the visual ($v$) and proprioceptive errors ($p$) according to the uncertainty of the sensory signal ($\sigma_v^2$ and $\sigma_p^2$), and the resultant aftereffect ($f(v,p)$) should be expressed as:

$$f(v,p) \sim \frac{\sigma_p^2}{\sigma_v^2 + \sigma_p^2} v + \frac{\sigma_v^2}{\sigma_v^2 + \sigma_p^2} p. \tag{S1}$$

If the uncertainty of the sensory signal (i.e., $\sigma_v^2$ and $\sigma_p^2$) is constant, $f(v,p)$ is linearly increased with the visual and proprioceptive errors. Fig. S1a indicates the how the dependence of the aftereffect on the visual error is modified by the additional proprioceptive errors. This pattern was apparently different from that observed by us (Fig. 3f-h).

**The modified version of the optimal estimation of the error**

The uncertainty of the signal could be increased with the signal intensity[3]. If the standard deviation of the signals linearly increases with the mean signal intensity, the aftereffect of the optimal estimation of the error can be represented as:

$$f(v,p) \sim \frac{\sigma_p^2 + k_p|p|}{\sigma_v^2 + k_v|v| + \sigma_p^2 + k_p|p|} v + \frac{\sigma_v^2 + k_v|v|}{\sigma_v^2 + k_v|v| + \sigma_p^2 + k_p|p|} p, \tag{S2}$$

where $\sigma_v^2$, $\sigma_p^2$, $k_v$, and $k_p$ are constants.

This model can reproduce the saturation of the aftereffect with the size of the error (Fig. S1b). However, the size of the modulation with the visual error becomes greater as the size of the proprioceptive error is increased. Furthermore, the aftereffect is linearly increased with the size of the error when the sizes of the visual and proprioceptive errors are identical (i.e., if $v = p = e$, then $f(v,p) \sim e$). These patterns are not consistent with the experimental results (Fig. 3f-h).

**Relevance of the error**

Wei and Kording[4] proposed a model in which the relevance of the (visual) error determines the aftereffect. Their model can be formulated as follows:

$$f(v,p) \sim \frac{N(v-p,\, \sigma^2)}{N(v-p,\, \sigma^2) + c} \left( \frac{\sigma_p^2}{\sigma_v^2 + \sigma_p^2} v + \frac{\sigma_v^2}{\sigma_v^2 + \sigma_p^2} p \right), \tag{S3}$$

where $N(v-p,\, \sigma^2)$ is the normal distribution function with mean $(v-p)$ and variance $(\sigma^2 = \sigma_v^2 + \sigma_p^2)$ and $c$ is a constant.

Fig. S1c demonstrates the pattern of aftereffects predicted by this model. This model nicely reproduced the saturation of the aftereffect with the error size. However, the aftereffect is linearly increased with the size of the error when the sizes of the visual and proprioceptive errors are identical (i.e., if $v = p = e$, then $f(v,p) \sim \frac{N(0,\, \sigma^2)}{N(0,\, \sigma^2) + c} e$), which was inconsistent with our results (Fig. 3f-h).

**Linear summation with decreased sensitivity to the error size**

Marko et al.[5] proposed a mechanism in which visual and proprioceptive memories are independently processed and then linearly integrated (summed). They also assumed that the sensitivity of each memory to the size of the error is decreased with the size of the error:

$$f(v,p) = \lambda_v \exp(-\tau_v |v|)\, v + \lambda_p \exp\left(-\tau_p |p|\right) p, \qquad \text{(S4)}$$

where $\lambda_v$, $\lambda_p$, $\tau_v$, and $\tau_p$ are positive constants.

Fig. S1d indicates the prediction of this model. Since the influence of the visual and proprioceptive errors is additive, the five curves in Fig. S1d are in parallel, which was again inconsistent with our experimental results (Fig. 3f-h).

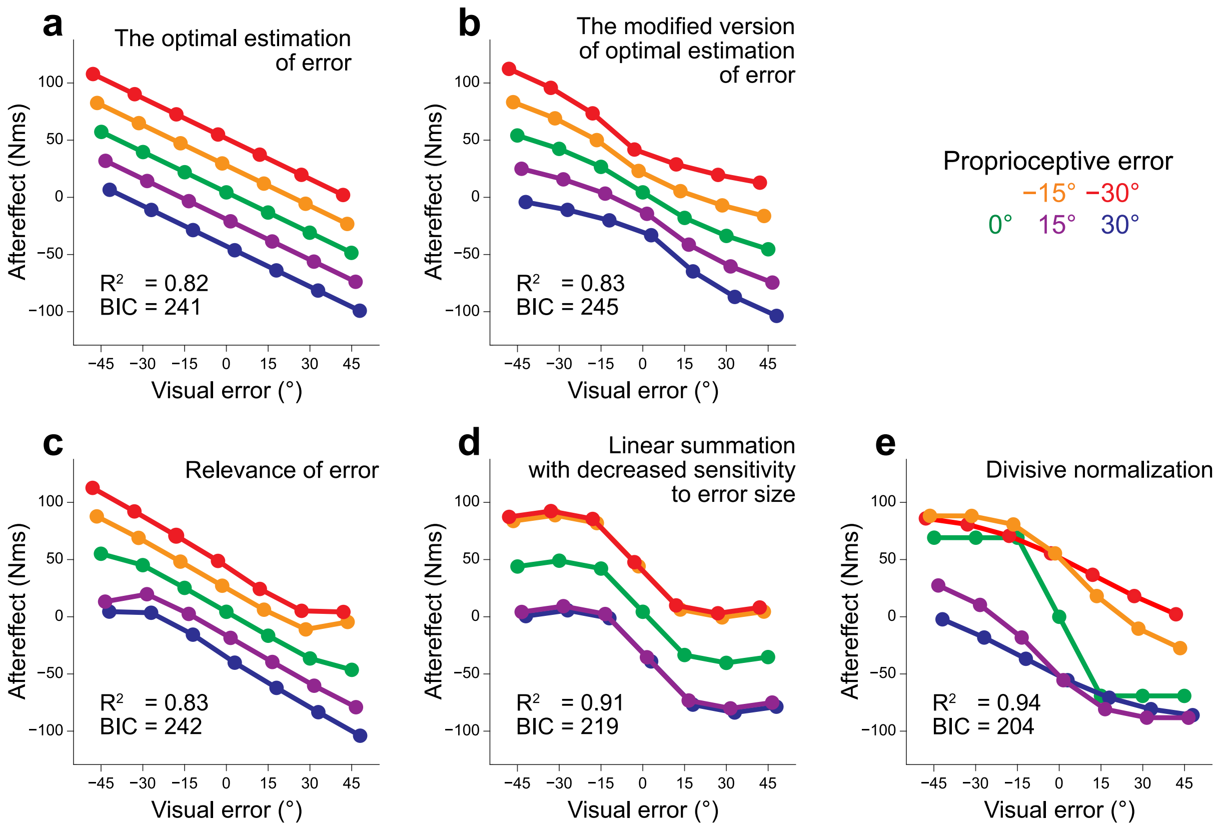


**Fig. S1. The results of data fitting by various computational models**

We fitted the results of Experiment 1 (Fig. 3F) with five different candidate models: **(a)** the optimal estimation of the error (Eq. S1) ($\sigma_v^2 = 289$, $\sigma_p^2 = 415$, gain = 3.21), **(b)** modified version of the optimal estimation of the error (Eq. S2) ($\sigma_v^2 = 13.9$, $\sigma_p^2 = 20.0$, $k_v = 0.47$, $k_p = 3.56 \times 10^{-10}$, gain = 3.43), **(c)** relevance of the error (Eq. S3) ($\sigma_v^2 = 9.00$, $\sigma_p^2 = 8.25$, $c = 1.93 \times 10^{-4}$, gain = 3.32), **(d)** linear summation with decreased sensitivity to the error size (Eq. S4) ($\lambda_v = 4.80$, $\lambda_p = 5.47$, $\tau_v = 3.51 \times 10^{-2}$, $\tau_p = 4.04 \times 10^{-2}$), and **(e)** our proposed models (Eq. 1) (same as Fig. 4A). Based on the R-squared and BIC comparison, we concluded that the divisive normalization model **(e)** is the most feasible model of multisensory integration for motor adaptation.

### 2. The state space model for modality-shared and modality-specific motor memory

Here, we attempted to explore the possibilities of how the motor memories could be updated in Experiment 3. A straightforward interpretation of the divisive normalization pattern of the aftereffect (Eq. 1 in the main text) is that the motor adaptation system updates the motor memory ($X$) according to the error information obtained from visual and proprioceptive perturbations ($v$ and $p$, respectively):

$$X(i+1) = \alpha X(i) + \frac{w_v v(i) + w_p p(i)}{\left\{\sigma^2 + k_v v^2(i) + k_p p^2(i)\right\}^{1/2}}, \tag{S5}$$

where $i$ represents the trial number and $\sigma$, $k$, and $w$ are constants (the subscripts represent visual ($v$) and proprioceptive ($p$) variables). Clearly, this model cannot explain the result of Experiment 3: The $X$ merely decayed during the washout phase. Therefore, the results of Experiment 3 indicate that vision and proprioception are likely to have distinct motor memories. One of the possibilities is that both visual ($x_v$) and proprioceptive ($x_p$) motor memory is updated by visual and proprioceptive perturbation ($v$ and $p$, respectively) and integrated according to the divisive normalization model.

$$x_v(i+1) = \alpha_v x_v(i) + \beta_v v(i), \tag{S6}$$

$$x_p(i+1) = \alpha_p x_p(i) + \beta_p p(i), \tag{S7}$$

$$X(i+1) = \frac{w_v x_v(i+1) + w_p x_p(i+1)}{\left\{\sigma^2 + k_v x_v^2(i+1) + k_p x_p^2(i+1)\right\}^{1/2}}, \tag{S8}$$

where $i$ represents the trial number, and $\alpha$, $\beta$, $\sigma$, $k$, and $w$ are constants (the subscripts represent visual ($v$) and proprioceptive ($p$) variables). However, this model has a critical flaw; even when both motor memories $x_v$ and $x_v$ increase with the motor adaptation, $X$ does not increase due to the constraint of Eq. S8. Therefore, the characteristics are unreasonable.

Another possibility is that each motor memory is updated by the divisively normalized error and the outputs are integrated as follows:

$$x_v(i+1) = \alpha_v x_v(i) + \frac{w_v v(i)}{\left\{\sigma^2 + k_v v^2(i) + k_p p^2(i)\right\}^{1/2}}, \tag{S9}$$

$$x_p(i+1) = \alpha_p x_p(i) + \frac{w_p p(i)}{\left\{\sigma^2 + k_v v^2(i) + k_p p^2(i)\right\}^{1/2}}, \tag{S10}$$

$$X(i+1) = x_v(i+1) + x_p(i+1). \tag{S11}$$

Note that this formulation is consistent with Eq. 1 in the main text. When the single trial adaptation is measured ($x_v(i) = x_p(i) = 0$), the aftereffect should be

$$X(i+1) = \frac{w_v v(i) + w_p p(i)}{\left\{\sigma^2 + k_v v^2(i) + k_p p^2(i)\right\}^{1/2}}, \tag{S12}$$

which is equivalent to Eq. 1 in the main text.

Fig. S2 indicates the result of the simulation by this state space model (Eqs. S9-S11). This model could reproduce the results of Experiment 3 (Fig. S2a) while reproducing the divisive normalization pattern in the single-trial adaptation (Experiment 1: Fig. S2b).

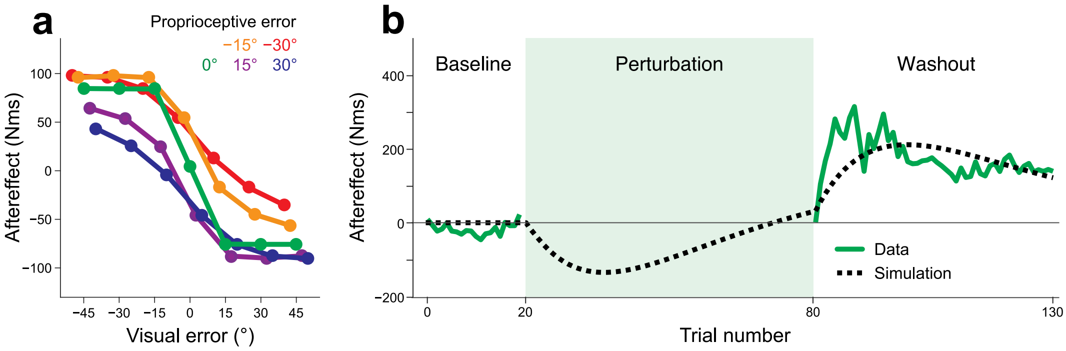


**Fig. S2. Simulation result of the modality-specific motor memory model**

(**a**) The aftereffect of single-trial adaptation (Experiment 1). (**b**) The aftereffect of visual and proprioceptive perturbations in the opposite directions (Experiment 3).

**Reference**

1. van Beers, R. J., Sittig, A. C. & Gon, J. J. Integration of proprioceptive and visual position-information: An experimentally supported model. *J. Neurophysiol.* **81,** 1355–64 (1999).

2. van Beers, R. J., Wolpert, D. M. & Haggard, P. When Feeling Is More Important Than Seeing in Sensorimotor Adaptation. *Curr. Biol.* **12,** 834–837 (2002).

3. Jones, K. E., Hamilton, A. F. & Wolpert, D. M. Sources of signal-dependent noise during isometric force production. *J. Neurophysiol.* **88,** 1533–44 (2002).

4. Wei, K. & Körding, K. Relevance of Error: What Drives Motor Adaptation? *J. Neurophysiol.* **101,** 655–664 (2009).

5. Marko, M. K., Haith, A. M., Harran, M. D. & Shadmehr, R. Sensitivity to prediction error in reach adaptation. *J. Neurophysiol.* **108,** 1752–63 (2012).
